# Ethylene impact on grapevine pistil temperature and fruit set

**DOI:** 10.1101/2022.08.08.502498

**Authors:** Christian Chervin, Olivier Geffroy

## Abstract

Ethylene is known to stimulate plant respiration, and this later is always associated with heat generation. The grapevine fruit set is dependent upon pistil temperature, controlling the pollen germination and ovule fertilization. This led us to test whether ethylene would be able to impact fruit set often limited in cool climate conditions, particularly in the case of *Vitis vinifera L*. cv. Malbec, a variety prone to poor fruit set. With this cultivar, we ran trials in growth chambers using the ethylene gas, and field trials using ethephon as an ethylene precursor. At 14.5 °C ambient temperature, we observed that ethylene at 1 and 10 ppm increased the stigma and pistil temperature to 15 °C, compared to 14.7 °C in controls. This minor temperature rise was associated with an increase of fruit set from 10 to 25 %. The field trials, conducted in the South West of France, confirmed this trend. Indeed, spraying a concentration as low as 216 mg/ha of ethephon onto Malbec at full bloom at 16 °C in the shade, led to an increase of fruit set from 35 % to 45 %. These experiments suggest that spraying ultra-low concentrations of ethylene precursors onto the grapevines prone to poor fruit set, during cool mornings, could improve fruit set and reduce crop losses.

## INTRODUCTION

The grapevine yield is obviously linked to flower physiology and good fruit set. For such development stages, the temperature is critical (Vasconcelos et al., 2009). These authors list three stages among others, which are particularly sensitive to variations in temperature with a critical impact on fruit set: (i) the pollen germination, (ii) the pollen tube growth within the style, and (iii) the fertilization of the ovule. The optimal temperatures for such stages are cultivar dependent, but an optimal range of 20 to 30 °C has been observed (Kozma, 2003). Small temperature rises, 15.5 to 17.5°C during the flowering time, may lead to major increases in bunch weight, 48 to 78 g (Vasconcelos et al., 2009). The grape cultivar Malbec, also named Cot, is known for having poor fruit set (Carrillo et al., 2020), with a fruit set around 20% with morning temperatures around 10°C and afternoon temperatures around 30°C.

The ethylene is a key plant hormone involved in fruit set (An et al., 2020), as it regulates plant pollination and fertilization processes. It finely modulates the pollen tube growth through modifications of cell wall and calcium gradient (Althiab-Almasaud et al., 2021). One aspect that has not been described in the two latter articles is the role of ethylene on organ temperature. In particular, it is well known that ethylene impairs the energy yield of respiration through the induction of the alternate oxidase pathway and uncoupling proteins (Wang et al., 2012; Hewitt and Dhingra, 2020), turning it into a thermogenic process (Moynihan et al., 1995). This respiration burst generating heat may be part of the plant response to chilling stress.

Thus, we studied whether ethylene was able to regulate fruit set by modulating the pistil temperature, in a controlled cool environment using fruiting cuttings. Then we tested whether the ethylene precursor, 2-chloroethylphosphonic acid, also called ethephon, was able to improve fruit set of Malbec clusters in a vineyard during cool mornings.

## MATERIALS AND METHODS

### 1. Growth chamber trials

#### 1.1 Growth chamber and fruiting cuttings set-up

The fruiting cuttings were generated as previously detailed (Ollat et al. 1998; Geny et al., 1998), using Malbec *(Vitis vinifera L.)* clone 595 woods collected at the experimental station of Institut Français de la Vigne et du Vin, Lisle sur Tarn (South West of France) in February 2019. Inflorescences on fruiting cuttings (Supplemental Figure S1A) were obtained by sequentially removing the first appearing leaves to limit the sink competition between vegetative and fruiting organs (Mullins and Rajasekaran, 1981; Ollat et al., 1998). One particularity of this plantlet model is that each fruiting bears one single inflorescence. The growing conditions were 17 °C/ 7 °C day/night for 16 h / 8 h, the day light intensity was 250 +/− 50 μmol m^-2^ s^-1^, the relative humidity was oscillating from 35 to 65 % (day/night). Batches of 6 plantlets were selected for the ethylene treatments as their flowers were between full bloom, stage 23, and 80% caps off, stage 25 (Coombe, 1995) (Supplemental Figure S1B).

#### 1.2. Ethylene treatments

The fruiting cuttings were incubated for 1 h at 14.5 °C in four crates as shown in Supplemental Figure S1C. The temperature was measured inside the crates using thermocouple thermometers (Voltcraft®, PL-120-T1, Conrad, France) over the treatment time. With the exception of the control treatment, ethylene gas was injected at 0.1, 1 and 10 ppm through the crate lid, using a hole performed with a hot needle. The ethylene concentration inside the crate was assayed by sampling gas just after closing, and before opening at the end of the incubation. The ethylene was measured by gas chromatography using a method previously described (Chen et al., 2020). A total of 40 Malbec fruiting cuttings were used in order to generate a set of data representing 4 batches of 10 inflorescences. The crates used for ethylene gassing contained 6 fruiting cuttings (Supplemental Figure S1C), the gassing experiments were repeated two times in order to treat 12 fruiting cuttings, one inflorescence by cutting. Crates were permutated to limit position bias. Over the remaining month of berry set and growth, some inflorescences showed development failure for all the studied treatments as a likely consequence of a bad rooting. Thus, we finally ended-up with 4 batches of 10 inflorescences, one batch per treatment.

#### 1.3. Measurements of pistil temperature

Six fruiting cuttings were randomly chosen among the 12 treated cuttings for each treatment, and were transferred in front of the camera within 30 min after the crate opening. Images of each inflorescence were taken using a A325sc thermal camera (FLIR Systems Inc., Croissy-Beaubourg, France) equipped with a close-up lens (1 × 100 μm, FLIR Systems Inc.) immediately after their removal from the crate. The camera had an angular field of view of 25° and a 320 × 240 microbolometer sensor, sensitive in the temperature range from −20 to 120 °C, with a thermal sensitivity below 0.05 °C. The distance between the camera and the pistil was 79 mm.

The images were analyzed in the following day using the FLIR ResearchIR software (Supplemental Figure S1D1 and D2). For each chosen inflorescence, three pistils were defined as regions of interest to determine average temperature and standard deviation, generating 6 batches of 3 pistils. Each batch corresponded to an inflorescence from a different fruiting cutting. The pistils were chosen according to the quality of picture focus.

The fruiting cuttings were then left in the growth chamber for one month, to later assess the fruit set.

#### 1.4. Fruit set assay

The fruit set is expressed as a percentage of flowers giving rise to a berry, for a chosen inflorescence. For the growth chamber trial, the number of flowers on one inflorescence were counted manually on pictures, before the treatment and the number of berries on one cluster were also counted manually on pictures 30 days after.

### 2. Field trials

#### 2.1 Vineyard and experimental set-up

Grapevine trials were set-up in the Cahors area (South West of France) in a private wine estate growing the Malbec cultivar, clone 598, onto the Riparia Gloire de Montpellier rootstock. The vines were 20 years old, the plantation density was 5000 vines per hectare. Six blocks of six vines were randomly spread over an experimental plot of 16 rows of 36 m each. One row was kept free of treatment between each treated row to prevent spray drift. The treatment took place in early June 2021, between full bloom and 80% caps off, as described in 1.1, with a morning temperature of 16°C in the shade. Five inflorescences were tagged (Supplemental Figure S1E) in each block, thus leading to 30 replicated measurements of fruit set per treatment. The inflorescence length was measured the day prior the treatments in order to estimate the number of flowers as explained in 2.2.

#### 2.2 Treatments and fruit set assay

Ethephon (Sierra^®^, Bayer SAS, Lyon, France) was sprayed on the vines at a rate of 110 L/ha, using a Cifarelli backpack mist blower (Supplemental Figure S1F). Ethephon was sprayed at three various final concentrations after dilution with water: 1.8, 18 and 180 mg/L. Controls were sprayed with water alone.

The percentage of fruit set was measured 30 days later. A regression between inflorescence length and flower number had been calculated using 25 inflorescences, as proposed by Ollat (1997), collected on a row adjacent to our experimental plot just before performing the treatments. At the treatment time, this enables to estimate the number of flowers by only measuring the length of all inflorescences. Then 30 days after the treatment, the clusters were sampled and the number of berries was determined manually.

### 3. Statistical analyses

Shapiro-Wilk and Brown-Forsythe tests, to estimate distribution normality and variance homogeneity, respectively, then Student or Welch t-tests to compare means, were performed using the Sigmaplot software v 14.5 (https://ritme.com/en/).

## RESULTS AND DISCUSSION

### 1. Ethylene treatments increased pistil temperature

As shown in Figure 1, the temperature of the pistils increased up to a maximal value of 15°C that was significantly higher than the control, after one hour of ethylene treatment at 1 ppm in a crate at 14.5°C. The other ethylene treatments also led to an increase in pistil temperature, although non-significant, but the trend is obvious. Thus, this shows that ethylene at very low concentrations can increase temperature of a plant organ. This may occur through the stimulation of the alternative oxidase pathway (Hewitt and Dhingra, 2020; Wang et al., 2012), but this was not demonstrated here, as our study mainly focused on the effect on fruit set. This could be a prospect for future studies. Even if the temperature increase due to 1 ppm of ethylene seems tiny: (+ 0.25°C in comparison to controls), it must be highlighted that this rise is the result of only one-hour incubation. And Vasconcelos et al. (2009) have shown that + 2°C at full bloom led to an increase in cluster weight by 40 %. The advantage of thermal camera was the non-destructive aspect of this measure, which allowed to keep the fruiting cuttings in the growth chamber to later observe the berries and calculate the fruit set. Additionally, it must be noted that the temperature rise following ethylene treatments lasted at least 30 min after crate opening, time to measure the pistil temperature of several inflorescences following a treatment batch. It would be interesting to further test for how long the pistil temperature rise is observable after treatment completion. Globally, each fruiting-cutting stood around two hours at 14.5°C before returning to the growth chamber. As additional experiments, various cold stress temperatures and durations could be tested.

**Figure 1:**
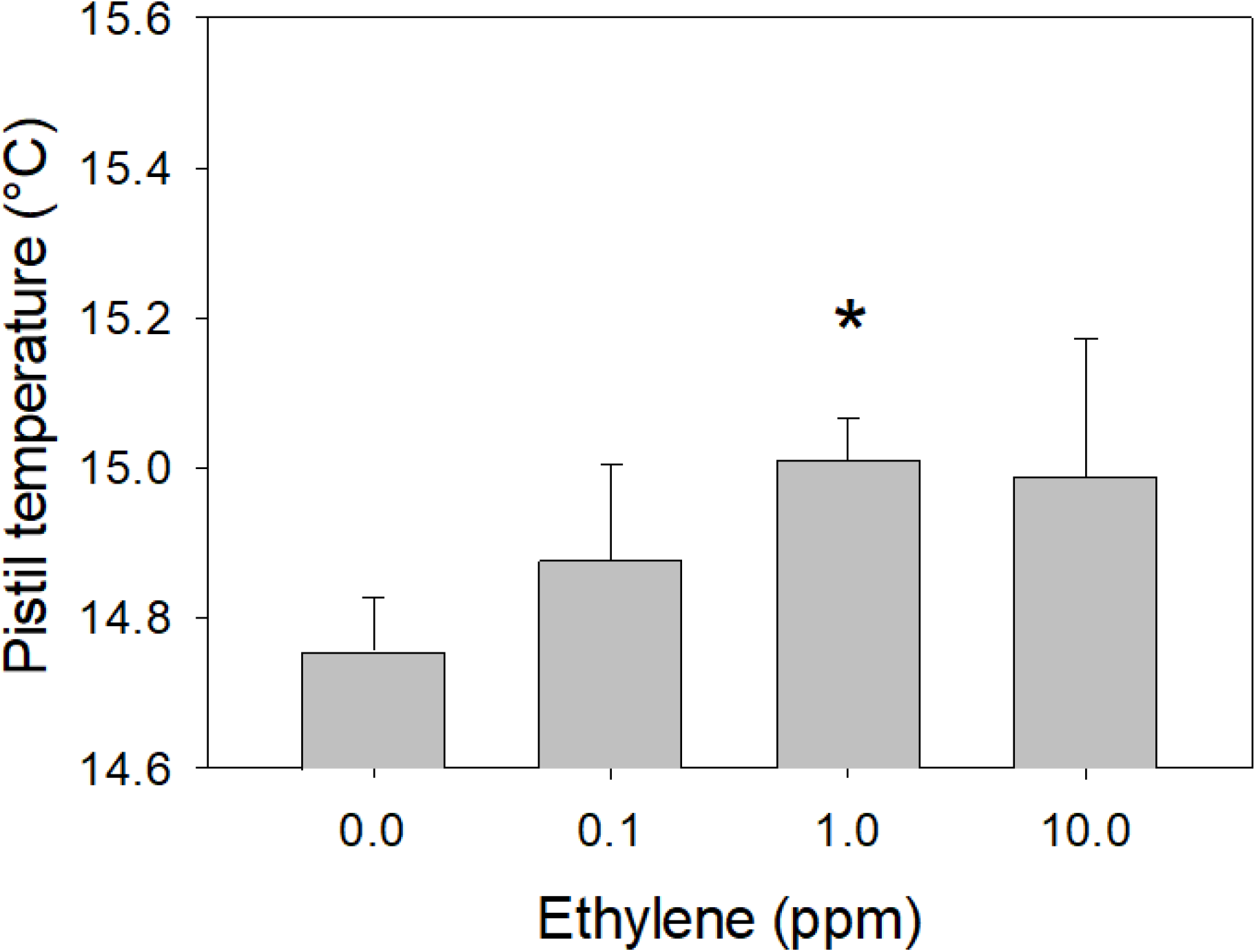
Changes in Malbec pistil temperature induced by exposure to exogenous ethylene at various concentrations in the growth chamber. n = 6 batches of 3 pistils, error bars show SE, and the * shows a significant difference between control and ethylene treated inflorescences by Student’s t-test at 5%.

### 2. Ethylene treatments increased fruit set

The Figure 2 shows that the two highest concentrations of ethylene, 1 and 10 ppm, generated a significant increase in fruit set from 8% to 25%. These percentages are quite low, for both control and ethylene treated inflorescences, but these are due to several factors. (i) Malbec is naturally a cultivar with low fruit set percentages (Carillo et al., 2020), (ii) we used cool growing conditions in order to induce poor fruit set, and (iii) fruiting cuttings are known to harbor a fruit set rate inferior to the rate observed in vineyards (Geny et al., 1998).

**Figure 2:**
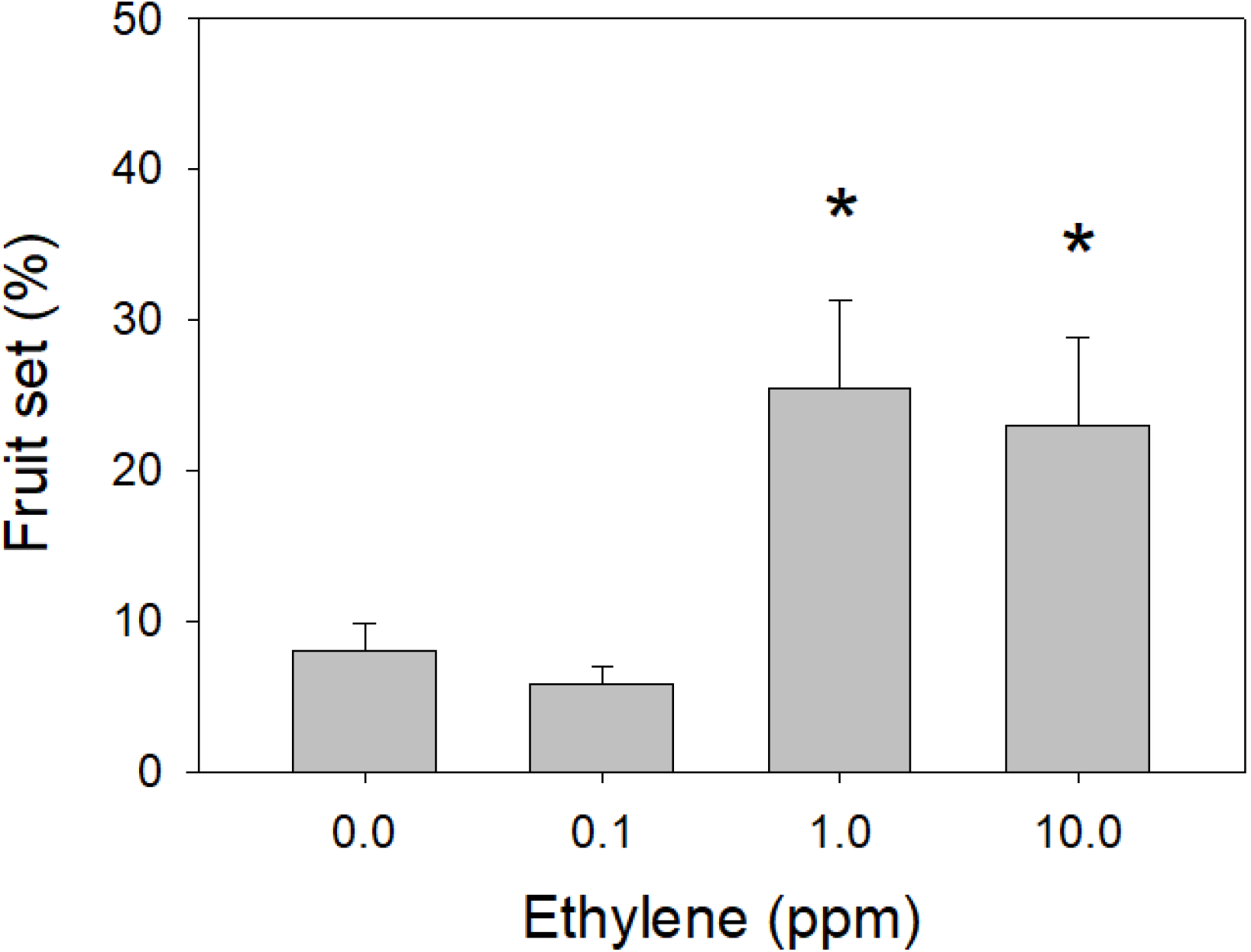
Changes in Malbec fruit set rate induced by exposure to exogenous ethylene at various concentrations in the growth chamber. n = 4 batches of 10 inflorescences, each inflorescence on a different fruiting cutting, error bars show SE, and the * shows a significant difference between control and ethylene treated cuttings by Welch’s t-test at 5%.

We cannot be certain that the three-fold increase in fruit set (Figure 2) is only related to the tiny increase in pistil temperature, as other metabolisms may be activated by ethylene (Althiab-Almasaud et al., 2021). For example, the ethylene effect on fruit set may involve some promotion of functional pollen maturation as observed by Völz et al. (2013) and/or increased fecundation (An et al., 2020). However, as previously shown, a small increase in temperature at flowering time can generate large increase in cluster weight due to better fruit set (Vasconcelos et al. (2009).

These observations were performed in the growth chamber, in a very controlled environment, because minimal pistil temperature shifts are easier to measure than in the field. In outdoor conditions, the thermal camera is harder to operate due to light pollution and wind, which impact image capture and temperature measurements.

### 3. Ultra-low concentrations of ethephon increased fruit set in the vineyard

During a preliminary set of experiments conducted in 2020 (unshown data), we observed that the optimal ethephon concentrations to test in the vineyard were between 1 and 200 mg/L in water.

The results obtained in the vineyard in 2021 are shown in Figure 3. An ultra-low concentration of 1.8 mg/L of ethephon led to a significant increase in fruit set rate from 35 % to 45 %. While increasing the ethephon concentration to 18 or 180 mg/L led to increased rate of fruit set too, but smaller than with 1.8 mg/L and non-significant. Such impact, although it remains weak in intensity, could lead to a yield enhancement, and have a positive effect on the economic balance of wine estates. This is particularly true given the expected low cost for this application. Ethephon is usually applied at a high concentration, approximately 1800 to 3600 mg/L, to perform chemical thinning (Weaver and Pool, 1971; Ferrara et al., 2016). The ultra-low concentration usage would allow to cover approximately 1000 hectares with a flask of 1 liter of commercial ethephon solution at 180 g/L of ethephon.

**Figure 3:**
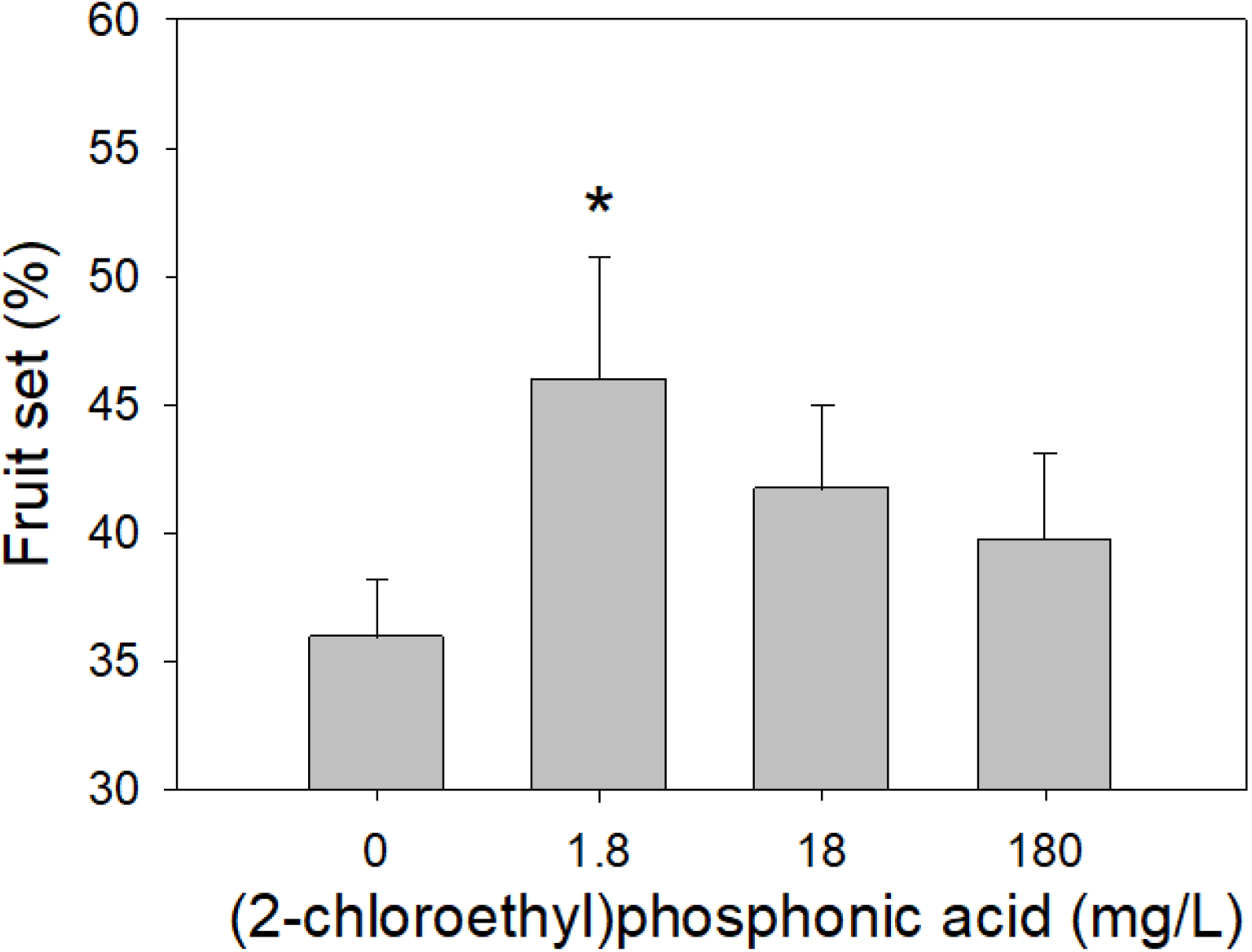
Changes in Malbec fruit set rate induced by spraying ethephon (2-chloroethylphosponic acid) at various concentrations in the field trial. n = 30 inflorescences, each inflorescence on a different grapevine, error bars show SE, and the * shows a significant difference between control and ethylene treated clusters by Welch’s t-test at 5%.

The fact that opposite effects are observed according to the applied ethephon concentration is not surprising. By “opposite” we mean that low concentrations improve fruit set, while high concentrations cause thinning. Such response is a classical “hormonal response” with stimulation of a metabolism at low concentration and inhibition of the same metabolism when the concentration increases (Weyers et al., 1995).

## CONCLUSIONS

We observed that ultra-low concentrations of ethylene, or ethylene precursor, can improve fruit set rate in cool conditions on Malbec, a cultivar usually leading to poor fruit set. There were several physiological evidences for this to happen: ethylene is known to boost alternative oxidase, alternative oxidase is thermogenic, and flower fertilization is temperature sensitive. However, the direct link between low ethylene concentrations and better fruit set rate had never been demonstrated. Additional experiments comparing a slight temperature increase to ethylene gassing could be an interesting test to separate effects by temperature and ethylene. And the critical period and duration of ethylene exposure could be further tested.

Practices such as girdling may improve fruit set (Tyagi et al., 2020 and references herein), but they are time consuming and not economically adapted for numerous vineyards. Ethylene at large concentrations is well known to produce chemical thinning (Weaver and Pool, 1971), but to our knowledge no research was conducted at ultra-low concentration.

As previously underlined, we did not decipher the whole process, as we firstly focused on the main output, that is improving fruit set, but, as outlined in discussion, the possible alternative targets of ethylene are numerous, from anther and pollen maturation, to pollen tube growth and pistil effects (An et al., 2019). Thus, decrypting the various metabolisms involved and testing ultra-low concentrations of ethephon with other cultivars remains an open area of research. Finally, our observations could lead to new development for ethephon at ultra-low concentrations.

## Acknowledgements

The authors are thankful to: Nathalie Ollat and Jean-Pierre Petit for their help to set-up the fruiting cuttings; Lauriane Heissat for the set-up of the growth chamber at the E.I. Purpan; Maxime Thorin, student at ENSAT, for running the fruit set trial in the vineyard, and Cédric Blanc, Domaine Lagrezette, for allowing the trials to be performed in the domain vineyard. Thanks to three anonymous reviewers for their time and fruitful suggestions.

---

**Supplemental Figure S1:**
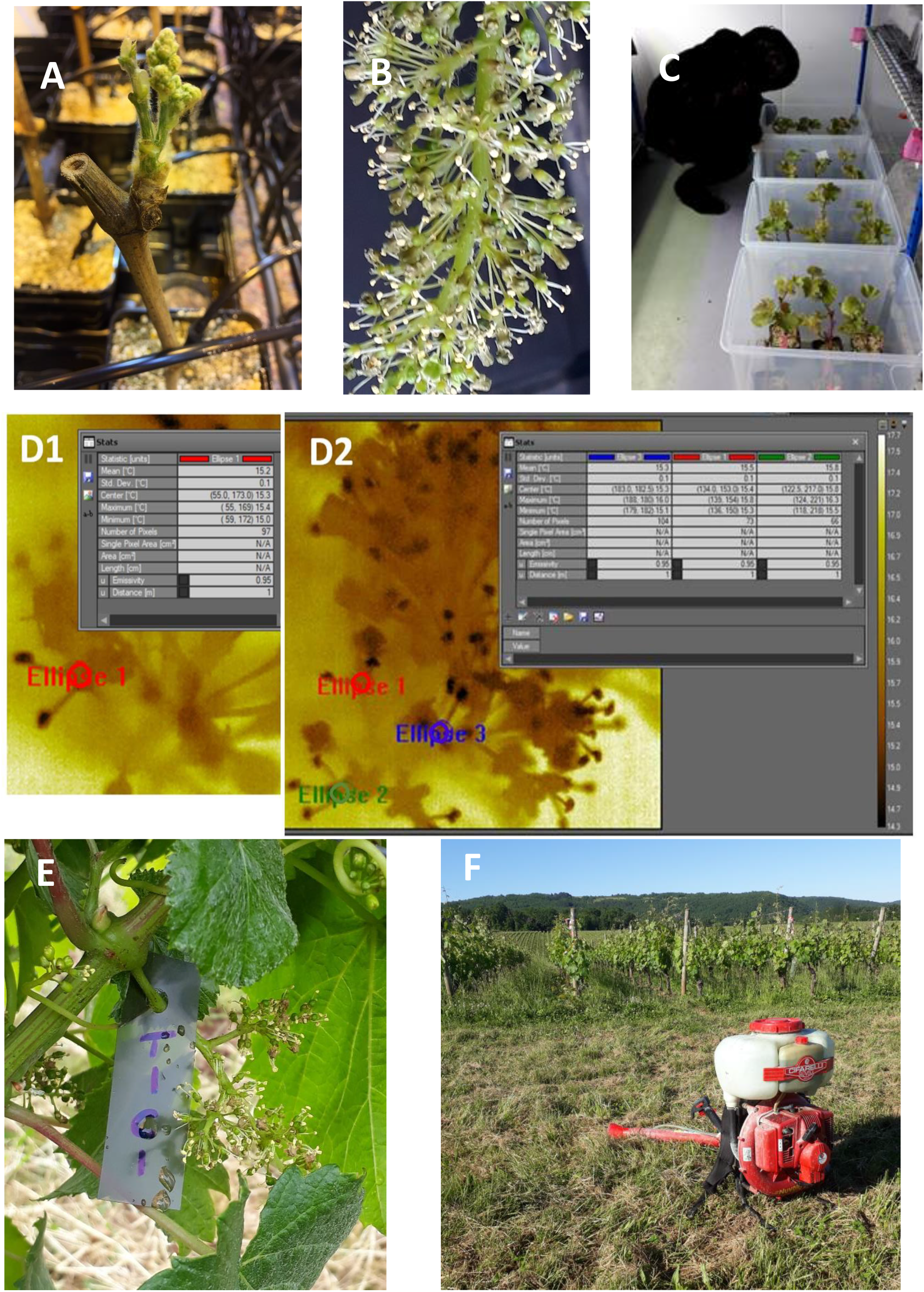
**A)** Inflorescence of a Malbec fruiting cutting; **B)** inflorescence between full bloom and 80% cap off for ethylene gas treatment, **C)** fruiting cuttings in crates for ethylene gas treatment, **D1 and D2)** IR images of pistils analyzed by the FLIR ResearchIR software, **E)** tagged inflorescence in the vineyard, **F)** mist blower used for spraying ethephon solution

